# Mechanical strain modulates Min patterning and division in *E. coli* filaments

**DOI:** 10.1101/2025.02.21.638242

**Authors:** Marta Nadal, Léna Guitou, Iago Díez, Juan Hurtado, Alejandro Martínez, Iago Grobas, Javier Buceta

**Author notes:** Contributing authors.

## Abstract

Bacteria often encounter physico-chemical stresses that disrupt division, leading to filamentation, where cells elongate without dividing. While this adaptive response enhances survival, it also exposes filaments to significant mechanical strain, raising questions about the mechanochemical feedback in bacterial systems. In this study, we investigate how mechanical strain influences the Min oscillatory system, a reaction-diffusion network central to division in *Escherichia coli*. Through a multidisciplinary approach combining quantitative fluorescence microscopy, biophysical modeling, microfluidics, and patterned growth substrates, we demonstrate that filamentous *E. coli* undergoes growth-induced buckling instability. This phenomenon alters the diffusivity of membrane proteins and modulates the spatiotemporal patterning of the Min system. Moreover, we show that this mechanochemical interplay determines division site positioning after stress relief, effectively creating a mechanical “memory” for cytokinesis. Our findings underscore the critical role of mechanical forces in bacterial filamentation and provide new insights into the functional implications of mechanobiology in microbial systems.

## 1 Introduction

Bacterial division has been extensively studied, typically under favorable growth conditions [1–3]. However, bacteria frequently face environmental stresses of diverse kind, both physical and chemical, that require a phenotypic adaptation [4–6]. Filamentation, a phenomenon where bacteria stop division but neither genetic replication nor elongation (i.e., growth), is a common adaptive response to various stressors, including antibiotic exposure [7–10], nutrient depletion [11], oxygen deficiency [12], and temperature changes [13]. The filamentation phenotype provides bacteria with advantages, such as colonizing heterogeneously adhesive surfaces [14] or facilitating cell-to-cell dissemination during host invasion [15]; but also poses challenges to bacterial cells. Thus, cells must cope with sizes up to hundreds of micrometers [16, 17] (in contrast to a size of a few microns when cells do not filament). As a consequence, bacterial filaments withstand significant mechanical stresses [18], compared to those in regular-sized cells, leading to an increased susceptibility to bending and buckling forces [19, 20].

The interplay between mechanical forces and signaling pathways, i.e., the mechanosensing properties, of bacteria during filamentation remains an unresolved question [21]. Notably, in filaments of rod-shaped bacteria, such as *E. coli* and *B. subtilis*, the cell wall growth rate has been shown to depend on mechanical stress and evidence suggests that MreB (an actin homologue) acts as a membrane curvature sensor [18, 22, 23]. However, how mechanical stimuli modulate other key signaling processes at the core of the filamentation phenomenon, such as the division machinery, is unclear.

The regulation of cell division has been extensively studied in *E. coli* [24–26]. In particular, the positioning of prospective septa, i.e., the so-called Z-rings [27, 28], is controlled by the Min oscillatory system (MinC, MinD, and MinE proteins) that follows a reaction-diffusion scheme [29, 30]. Thus, when *E. coli* undergoes filamentation, the Min system exhibits a standing-wave, multinode, pattern, whose potential division sites are located at the minima of the time-averaged Min signal [27, 31, 32]. Interestingly, experiments on deformed spheroplasts [33], in *E. coli* with aberrant shapes [34, 35], and in patterned substrates [36], suggest that MinD localizes in regions of high curvature. While the molecular mechanism underlying this phenomenon is, to the best of our knowledge, still under debate, a plausible reason is the reported positive feedback loop between the local membrane curvature and protein concentration [37]. These facts raise the intriguing question of whether or not in bacterial filaments, prone to bend and buckle, Min patterning is influenced by mechanical stress (or the other way around). Moreover, given that Min patterning determines the prospective locations for cell cleavage, does the mechanical stress condition the division sites in filaments when the environmental stress ends and cytokinesis resumes?

Herein, we address these questions using a multidisciplinary approach that combines quantitative fluorescent time-lapse microscopy, micro-patterning of growth substrates, biophysical models, Fluorescence Recovery After Photobleaching (FRAP) analyses, and microfluidics. Our results reveal that bacterial filaments undergo growth-induced buckling instability that leads to a mechanochemical interplay between the Min oscillatory system and the mechanical strain. Further, we explore the post-stress consequences of such an interplay and conclude that it sets preferential division sites. Altogether, our study sheds light on the mechanobiology of bacterial filaments and its functional role.

## 2 Results

### 2.1 *E. coli* growing filaments undergo a buckling instability

In our experiments we used the antibiotic aztreonam to induce *E. coli* filamentation. Aztreonam inhibits the cell division protein FtsI that catalyzes cross-linking of the peptidoglycan cell wall at the septum [38]. At low, sub-lethal, concentration values, aztreonam does not compromise the cell membrane and the Z-ring assembles, but division is hindered, thus leading to the formation of long bacterial filaments [9, 39]. In our case, *E. coli* cultures were grown in M9 minimal medium to an OD_600_ of 0.3 before inoculation onto 2% agarose pads containing 10 *µ*g/ml of aztreonam (Methods). Under these conditions, time-lapse microscopy revealed cell filamentation (Fig. 1A, Movie 1). Growing filaments exhibited a buckling instability which became more likely for filaments longer than ∼10 *µ*m (Fig. A1). This instability is, at first, counterintuitive, since cell elongation could be thought as a tensile stress whereas buckling develops due to compressive stresses [37]. In that regard, our estimation of the critical load required to induce filament buckling is on the order of 𝒪(10^1^) nN (Methods). Buckling led to high curvature values with radii of curvature smaller than the characteristic length of non-filamentous *E. coli* cells: |*κ*| *>* 0.5 *µ*m^−1^ (Fig. 1A). In this context, we used |*κ*| as a proxy for mechanical strain.

**Fig. 1.**
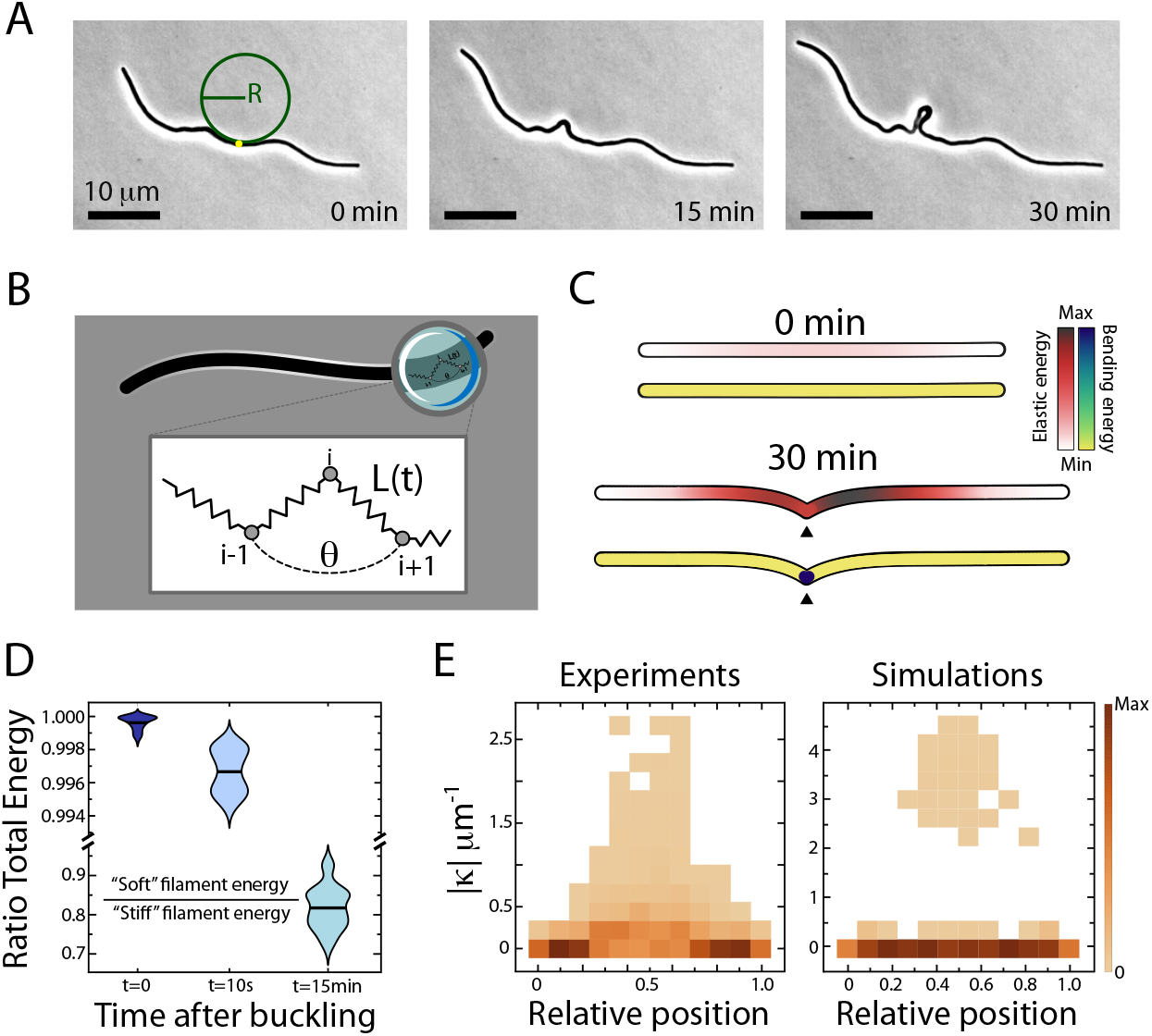
Growing *E. coli* filaments undergo a buckling instability. **A**. Phase contrast images of a representative *E. coli* filament growing on 10 *µ*g/ml aztreonam agarose pads. Following cell segmentation and tracking (Methods), the curvature of the filaments is computed along the midline. At a given location (e.g. yellow point) the curvature, |*κ*|, is computed as the inverse of the radius of the circumference that is tangent to that point (green circle): |*κ*| = 1*/R*. As time progresses and the filament grows, the snapshots reveal a buckling instability where large values of |*κ*| are reached by the central region of the filament (see **E**). **B**. Cartoon of the mechanical toy model. Forces are exerted by a chain of connected springs (black). Nodes (grey circles) are subjected to elastic and bending forces. Due to cellular growth, the elastic energy relies on the time-dependent distance, *L*(*t*), between consecutive nodes (Methods). Bending energy depends on the angle *θ* as determined by neighboring segments. **C**. Snapshots of a simulated growing filament (initial length: 50 *µ*m; doubling time: 50 min). Color scales for each time point indicate the elastic (top) and bending (bottom) energies using pair-wise moving average coordinates (Methods). As time progresses elastic energy accumulates by the central region. Buckling leads to a relaxation of the elastic energy at the expense of an increase of the bending energy (black triangles). **D**. Ratio of the total energy between soft and stiff filaments (soft: 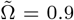; stiff: 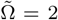 sample size: *n* = 100 simulations). As time progresses the panel shows that stiff filaments that are not able to buckle increase their energy. **E**. Density histograms of the curvature of filaments as a function of the relative position as obtained in experiments (left) and simulations (pair-wise moving average coordinates; right). Sample size: *n* = 22 (experiments) and *n* = 100 (simulations). The color scale represents probability. Larger curvatures (buckling) accumulate by the central region of the filament.

To understand this phenomenon we developed a mechanical toy model. The model consists in a 2D chain of massless nodes connected by linear springs. Segments connecting nodes are also subjected to bending forces (Methods, Fig. 1B). The springs were assumed to be identical (i.e., same elastic and bending constants). In the model we neglected inertial effects and included energy dissipation. Thus, the model comprises the minimal components required to mimic the filament mechanics at low Reynolds numbers by disregarding the 3D structure of the cells, the adhesion forces to a substrate, and plastic deformation effects (see Discussion).

The theoretical analysis and our simulations confirmed that, as expected, if the equilibrium length of the springs, *l*_0_, is kept constant, then the simulated filaments reach a straight configuration regardless of the initial condition (Methods). However, if growth is included such that the equilibrium length depends on time, i.e. *l*_0_ (*t*), then buckling develops (Movie 2). Consequently, our model argues in favor of cellular growth as the driven force underlying buckling. On the one hand, the growth of the springs located at the middle of the chain is dampened by the growth of the surrounding springs that push back. On the other hand, springs located by the poles of the chain find less opposition against growth. As a result, elastic energy accumulates at the central region of filaments (compressive stress) and energy relaxation can only be achieved by buckling. This buckling dynamics is revealed in our model by bending events at different length scales. Thus, local stresses at a scale of few nodes cause zigzag configurations in the chain of springs. In addition, at a larger length scale, there is macroscopic buckling as revealed by the changes of the local curvature of the midline as experimentally shown (e.g. Fig. 1A). In order to distinguish between these different, yet complementary, phenomena, and properly compare simulations and experiments, we implemented a moving average approach (Methods).

As shown in Fig. 1C our model reproduces that the accumulation of elastic energy (compressive stresses) due to growth decreases at the expense of an increase of the bending energy. The total energy decrease due to the buckling instability can be observed in Fig. 1D by comparing filaments with different (dimensionless) bending moduli: 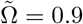 (“soft” filament) vs. 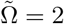 (“stiff” filament). Thus, as time progresses, the simulations show that stiff filaments that are not able to buckle heap up energy with respect to soft filaments. The model also predicts that since compressive stresses accumulate at the middle of the filament, then buckling is more likely to develop at that location. In that regard, Fig. 1E shows that, both in experiments and simulations, the region of the filament with the largest curvature is the central one. The experimental Fig. 1E also allows us to estimate the onset of buckling for |*κ*| *>* 0.5 *µ*m^−1^ as these values are reached only at the central region of the filament.

In summary, the mechanical model explains the buckling phenomenon observed in growing *E. coli* filaments. Buckling events occur more likely by the central region because the filament ends are able to elongate freely, whereas the middle section must exert larger forces in order to push and displace the rest of the filament. This mismatch between the growing dynamics of different regions of the filament leads to an accumulation of compressive stresses that ultimately cause the filaments to buckle.

### 2.2 Min patterning is modulated by the local curvature of filaments

In order to study how mechanical stress affects signaling proteins related to the filamentation phenomenon, we focused on the Min system. The MinCDE oscillatory system prevents FtsZ polymerization and, consequently, plays an important role in the filamentation and cell division processes [40–42]. In particular, MinD protein localizes either at the cell membrane or in the cytosol depending on its ATP activity and it is a proxy for the activity and localization of MinC and MinE. In our study, we used an *E. coli* strain that expresses the recombinant fluorescent protein YFP::MinD (Methods). We tracked and measured, simultaneously, MinD and the curvature of the middle cell axis of *E. coli* filaments by quantitative time-lapse microscopy (Fig. 2, see also Movie 3). In our analyses, we focused on growing filaments where MinD displayed a standing wave pattern. Additionally, this allowed us to pinpoint the location of the non-functional FtsZ septa during filamentation (regions with a sustained low concentration of MinD). Interestingly, we observed that MinD concentration (i.e. the amplitude of the pattern) was dependent on the level of curvature, Fig. 2A. In particular, the average intensity of the MinD signal seemed to increase in buckled regions and, concurrently, the imaging data suggested that the nodes of the MinD pattern (i.e. local minima) tallied with large curvature peaks.

**Fig. 2.**
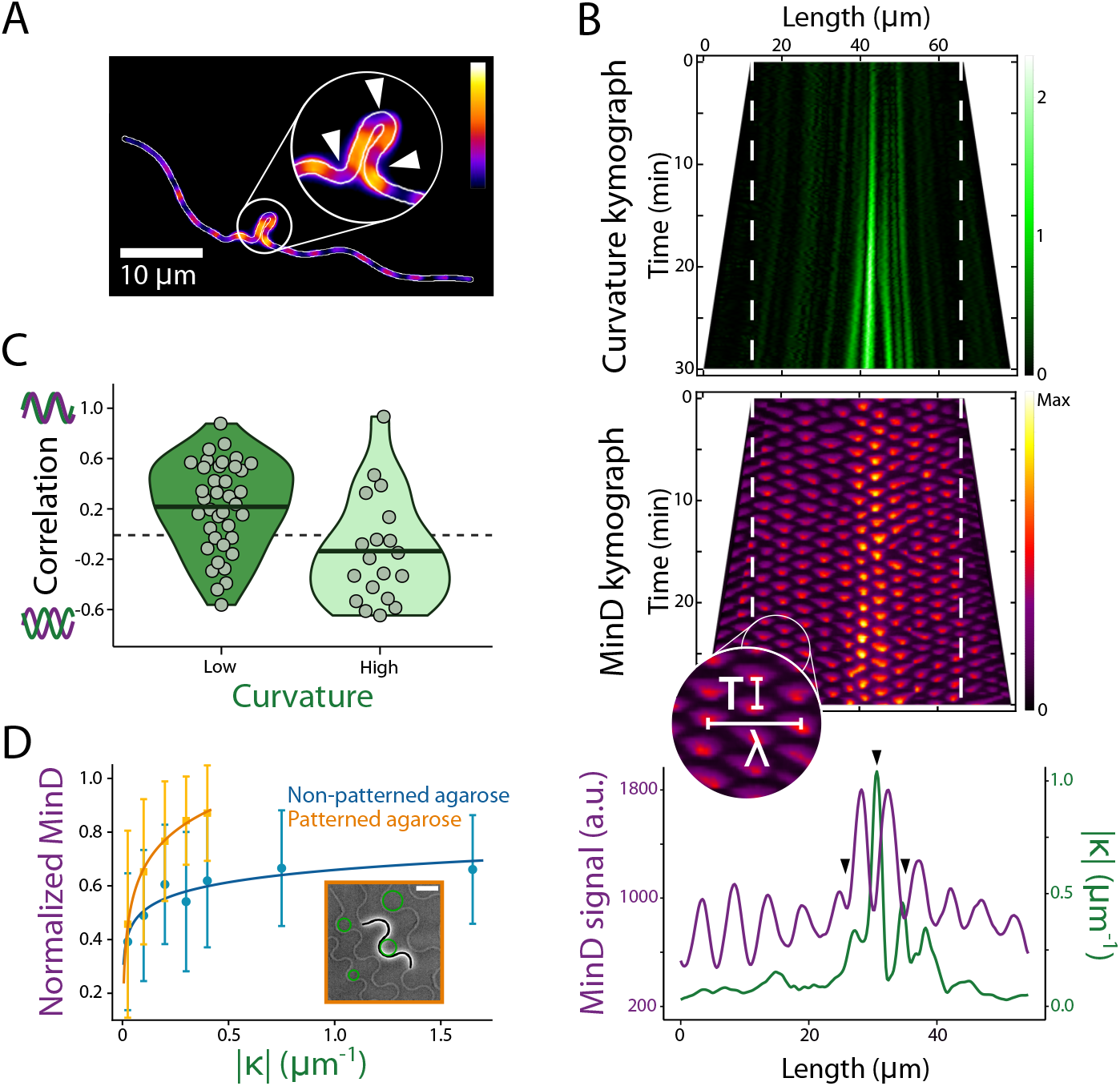
Min patterning is modulated by the local curvature of filaments. **A**. Fluorescence channel (YFP, false color scale) of the illustrative filament shown in Fig. 1A (*t* = 30 min). The zoomed region (circle) highlights the increase of MinD in buckled regions. In addition, the white arrows indicate the anti-correlation effect between MinD and locations with a high curvature. **B**. Top: Kymographs of the filament shown in panel **A** representing curvature (top) and MinD intensity (middle) profiles along the filament middle axis as a function of time. Color scales account for the curvature (*µ*m^−1^) and for MinD concentration (arbitrary units). The zoomed region in the MinD kymograph indicates the period, *T*, and the wavelength, *λ*, of the MinD standing-wave pattern. Bottom: Overlay of the time average of MinD and curvature profiles. Black arrows highlight the locations pinpointed in panel **A** by means of white arrows. Averages are computed within the region delimited by the white dashed lines. **C**. Violin plots showing the distribution of values of the Pearson’s correlation coefficient between MinD and curvature (sample size: *n* = 19 filaments where high and low curvature regions were analyzed separately, Fig. A2C, Methods). Black solid lines represent the mean (Low: 0.22; High: −0.14). **D**. Normalized MinD intensity (mean and standard deviation) as a function of the curvature in non-patterned agarose pads (same dataset as in **C**) and in micro-patterned growth substrates (*n* = 20 filaments). Solid lines are, in both cases, a logarithmic fit: Eqn. (1), Methods. Inset: Illustrative phase contrast image of an *E. coli* filament growing in a micro-patterned agarose pad (scale bar: 10 *µ*m).

To quantitatively assess the interplay between MinD patterning and buckling, we computed kymographs and performed temporal averages of MinD and curvature profiles (Fig. 2B). As expected, the number of nodes of the standing wave increased as the filament grew but properties of the pattern, such as the wavelength (at any given time) and the period (at any given location), were not affected by buckling (Fig. A2A). Further, these values were fairly conserved among filaments: *λ* = (7.6 ± 1.0) *µ*m and *T* = (55 ± 10) s (Fig. A2A), in agreement with previous studies [27]. Our analysis confirmed that MinD peaks avoided regions with high curvature (|*κ*| *>* 0.5 *µ*m^−1^). To deepen into this effect, we computed the Pearson’s correlation coefficient between the averaged MinD and the curvature profiles (Methods) and differentiate between regions with high (|*κ*| *>* 0.5 *µ*m^−1^) and low (|*κ*| *<* 0.5 *µ*m^−1^) curvature values (Fig. A2B). The results, Fig. 2C, corroborated a significant trend towards a negative correlation in the high curvature regions and towards a positive correlation in the low curvature regions (Mann-Whitney *U* test: *α* = 5·10^−2^ significance level). As for the quantitative relationship between MinD concentration and buckling, we computed the value of the normalized intensity (per filament) of the wave envelope as a function of the curvature, Fig. 2D (Methods, Fig. A2D). The analysis confirmed an increase of MinD as a function of the curvature that reaches a plateau for curvature values above 0.5 *µ*m^−1^ (Fig. 2D non-patterned agarose).

Given that our results indicate that buckling instability naturally develops in the central region of filaments (Fig. 1E), we further tested whether MinD accumulation preferentially occurs at regions of higher local curvature versus at locations by the middle (where growth-induced compressive stresses accumulate). To that end, we designed micro-patterned growth substrates (Methods, Fig. A3), where we externally control the bending and curvature of the filaments along their body length (inset Fig. 2D and Movie 4). Our analysis of MinD intensity as a function of the curvature revealed in this case a similar trend (Fig. 2D: patterned agarose). We notice that micro-patterned pads could not be used to perform a MinD-curvature correlation study (i.e., Fig. 2C). As mentioned above, that analysis requires to achieve local curvatures above 0.5 *µ*m^−1^. However, we observed that *E. coli* filaments would escape the regions of the guiding growth patterns with curvatures larger than 0.4 *µ*m^−1^. Therefore, our data supports that MinD accumulation is driven by local curvature rather than by growth-induced mechanical strain, and, overall, the existence of a modulation of Min patterning due to curvature effects. These findings align with previous studies conducted on non-filamentous *E. coli* cells [37] (Discussion).

### 2.3 MinD pattern modulation is driven by a diffusion drop in buckled regions

Previous studies have shown that membrane curvature and mechanical stress alter the assembly dynamics of proteins as well as their diffusive properties [37, 43–47]. In that regard, since local changes in protein diffusion lead to local modifications in protein concentration, we wondered whether the modulation of the MinD pattern in buckled regions is driven by this physical phenomenon. To test this hypothesis, we performed Fluorescence Recovery After Photobleaching (FRAP) assays and numerical simulations. Given the oscillatory behavior of MinD, we devised a novel analysis methodology of FRAP data that weights up the standing-wave patterning properties of the signal (Methods, Fig. 3A). Our analysis relies on the fact that a standing wave can be decomposed as the sum of two waves traveling in opposite directions with phase velocities *c* = *λ/T*. To characterize the effect of buckling in protein diffusivity, we photobleached MinD and computed the dimensionless ratio between the fluorescence recovery speed, *v*_*r*_, and *c*. Here, *v*_*r*_ is defined as *v*_*r*_ = *d/t*_*r*_, where *d* is the distance from a MinD peak in the non-photobleached domain (emitter) to the location where the recovery (≥ 30%) of the MinD signal is observed (receiver), and *t*_*r*_ is the recovery time following photobleaching (Methods, Fig. 3B). We note that the normalization of *v*_*r*_ with respect to *c* accounts for possible differences (i.e. variability) among filaments with respect to their wave properties. Consequently, the larger *v*_*r*_*/c*, the faster the signal propagation due to the transport mechanism underlying patterning, i.e. diffusion.

**Fig. 3.**
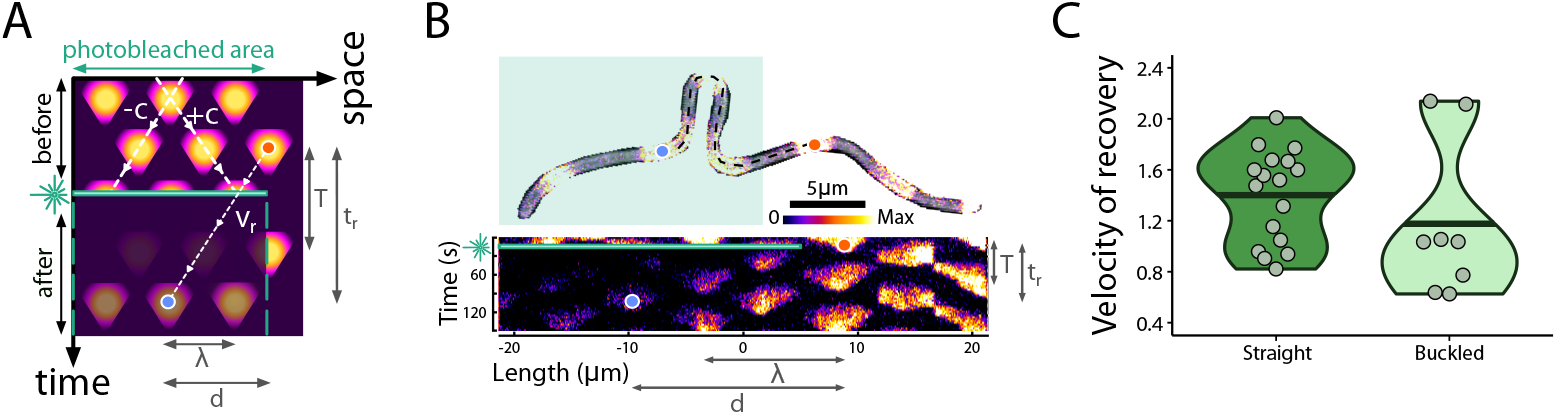
FRAP experiments suggest a decrease in the recovery velocity of MinD pulses in buckled filaments. **A**. Cartoon illustrating the FRAP experimental quantification (see text). **B**. Top: Composite of bright-field and MinD fluorescence of a filament before photobleaching. The colored area highlights the photobleached region. The blue circle indicates the pulse for which the fluorescence recovery is analyzed (receiver). The orange circle highlights the non-photobleached pulse before the buckled region (emitter). Bottom: Kymograph of the FRAP experiment performed over the filament shown in the top. **C**. Violin plots showing the dimensionless velocity recovery, *v*_*r*_*/c*, of the pulse’s recovery for straight (*n* = 17) and buckled (*n* = 8) filaments. Black lines account for the mean (Straight: 1.40; Buckled: 1.18). Buckled filaments reveal a larger recovery thus indicating a slower diffusive transport.

To test this approach, we performed several control experiments (Fig. A4). In the case of non-filamentous bacteria showing one pole-to-pole MinD pulse, photobleaching the pulse precludes recovery within the temporal observation window (∼2.5 ×*T*, Fig. A4A). However, photobleaching one cell pole while there was a signal outside the photobleached region led to MinD recovery within the observation window (Fig. A4B). This hints that signal recovery requires MinD traveling from a non-photobleached region within the filament. To further investigate this, we conducted the same experiment using short filaments. Consistent with the previous results, when we photobleached all of the MinD (Fig. A4C), we observed no signal recovery. However, when we photobleached one of the pulses, while leaving at least one other MinD pulse intact, recovery was detected (Fig. A4D). We applied the aforementioned methodology to buckled (|*κ*|_*max*_ *>* 0.5 *µ*m^−1^) and straight (|*κ*|_*max*_ *<* 0.5 *µ*m^−1^) filaments. Our results, Fig. 3B, indicate that buckling slows down the velocity recovery, *v*_*r*_*/c*, therefore suggesting a decrease in protein diffusivity in curved regions as previously shown [45–49].

To strengthen our findings from the FRAP experiments, and given that our experimental setup cannot distinguish between the fluorescence signals of membrane-bound and cytoplasmic MinD, we conducted numerical simulations. In that regard, since the cross-sectional shape of the bacterial filaments remains unchanged due to bending/buckling, we do not expect that the transport properties of MinD in the cytoplasm are affected [50]. In contrast, the curvature of membranes has been shown to influence protein diffusivity [48, 51]. As a result, we anticipated that the diffusion of membrane-bound MinD is reduced.

Our approach is based on a model [52] that, while rudimentary from a signaling viewpoint (e.g. MinC is excluded), accounts for the essential elements required to produce a Min standing-wave pattern (Fig. 4A, Methods). Specifically, the model includes the dynamics of both membrane-bound and cytoplasmic forms of MinD and MinE.

**Fig. 4.**
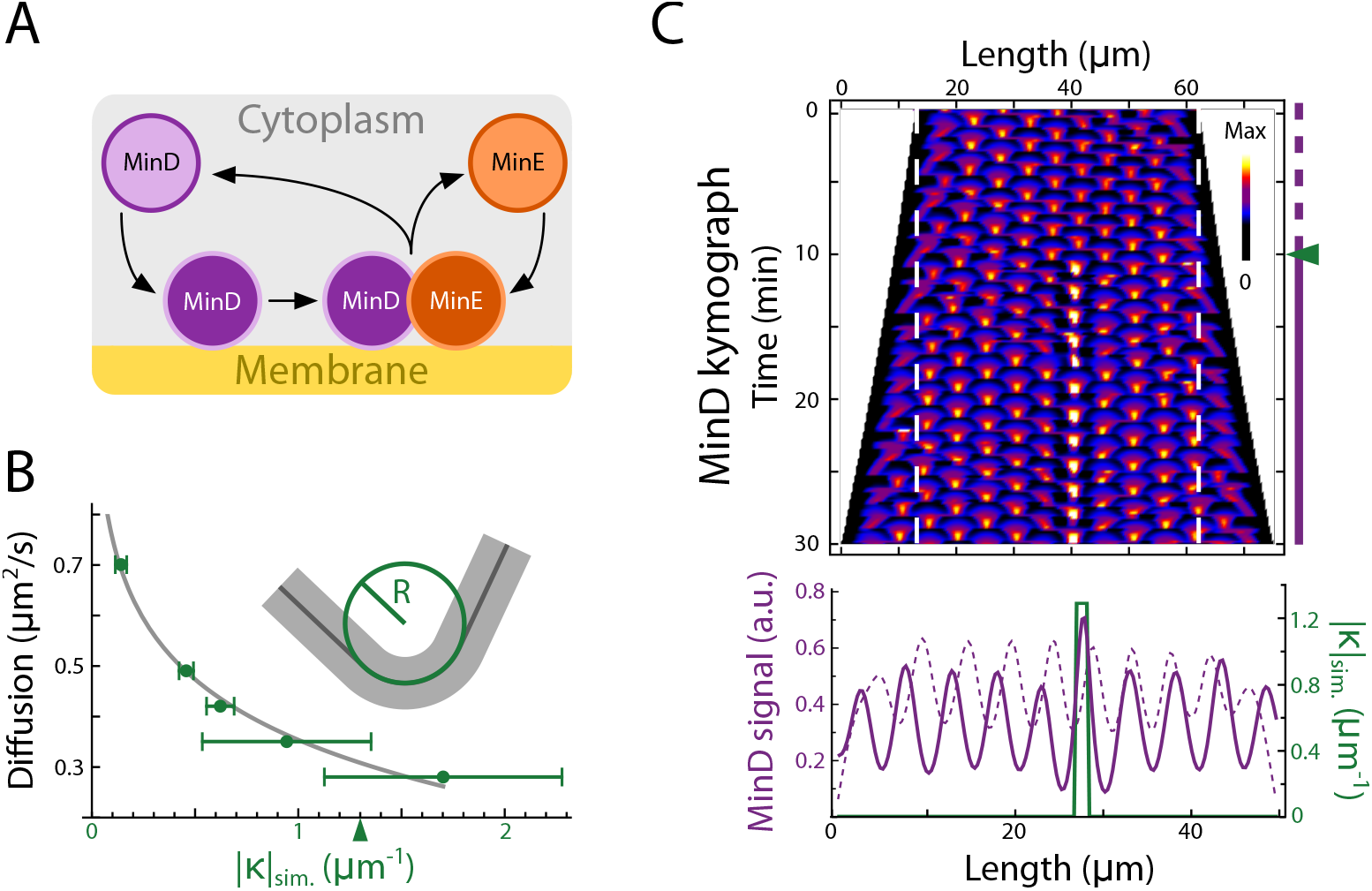
MinD signaling modeling supports the hypothesis of a membrane diffusion drop due to buckling. **A**. Cartoon of the simplified Min oscillatory system model [52]. **B**. MinD membrane diffusion, 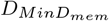, as a function of the curvature |*κ*|_*sim*._ obtained in calibration simulations (Methods, Eqn. 4). The solid line represents the fit to Eqn. 5. The inset represents the buckled region of the filament used in the simulation shown in panel **C** with a curvature |*κ*|_*sim*._ = 1*/R* = 1.3 *µ*m^−1^ (green arrow). **C**. Spatio-temporal evolution of *MinD*_*sim*._ = *MinD*_*mem*._ + *MinD*_*cyto*._ obtained in simulations of a growing filament. Top: Kymograph of a filament with an initial length of 50 *µ*m growing for 30 min (doubling time: 50 min). For *t* ∈ (0, 10) min (dashed purple line) no buckling is implemented: 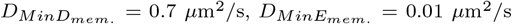. For *t* ∈ (10, 30) min (solid purple line), the diffusion coefficients of proteins are set to a 40% of their values within a 1.5 *µ*m-wide band: 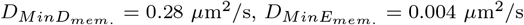. Bottom: Temporal averages of *MinD*_*sim*._ for *t* ∈ (0, 10) min (dashed purple line) and for *t* ∈ (10, 30) min (solid purple line) within the region delimited by the white dashed lines shown in the kymograph. The green line represents the curvature profile of the simulated filament, |*κ*|_*sim*._.

In our simulations we implemented an exponential growth of the filaments (i.e. elongation) and a reduction in diffusion in a 1.5 *µ*m region —approximately the width of a MinD peak. We calibrated the value of the diffusion as a function of the level of curvature using experimental data (Fig. 4B, Fig. A5A, Methods). Thus, we estimated that, for example, a buckled filament configuration with a curvature of approximately |*κ*| ∼1.5 *µ*m^−1^ would result in a membrane protein diffusivity reaching ∼40% of its original value. Under those conditions, the MinD accumulation pattern in buckled regions observed in our simulations closely resembled the experimental results (cf. Fig. 2B middle and Fig. 4C). Notably, when the same diffusion drop was applied just at the cytoplasmic level, the characteristic accumulation patterning observed in experiments was not reproduced (Fig. A5B). In summary, FRAP experiments and numerical simulations support the idea that buckling modifies the membrane diffusivity of Min proteins, which in turn causes protein accumulation.

### 2.4 Post-stress filaments preferentially divide in regions of high curvature

Herein we have shown that curvature peaks positively correlate with sustained MinD signal minima (i.e., as shown in Fig. 2C, high curvature —indicating large mechanical strain— and MinD maxima are anti-correlated). This observation prompted us to investigate whether Min patterning and mechanical effects might synergistically determine preferred division sites in post-stress conditions.

To verify this, we conducted microfluidic experiments adapting the protocol established by Wehrens *et al*. [32] (Methods). In these experiments, cells growing and dividing in M9 medium were exposed to a stressor medium (M9+aztreonam) to induce filamentation over a 3 − 5 h period, during which the cells elongated and subsequently 10 buckled. To resume cytokinesis, the cells were exposed again to just M9 medium and the bacterial filaments started to divide [16, 32, 53] (Fig. 5A and Movie 5).

**Fig. 5.**
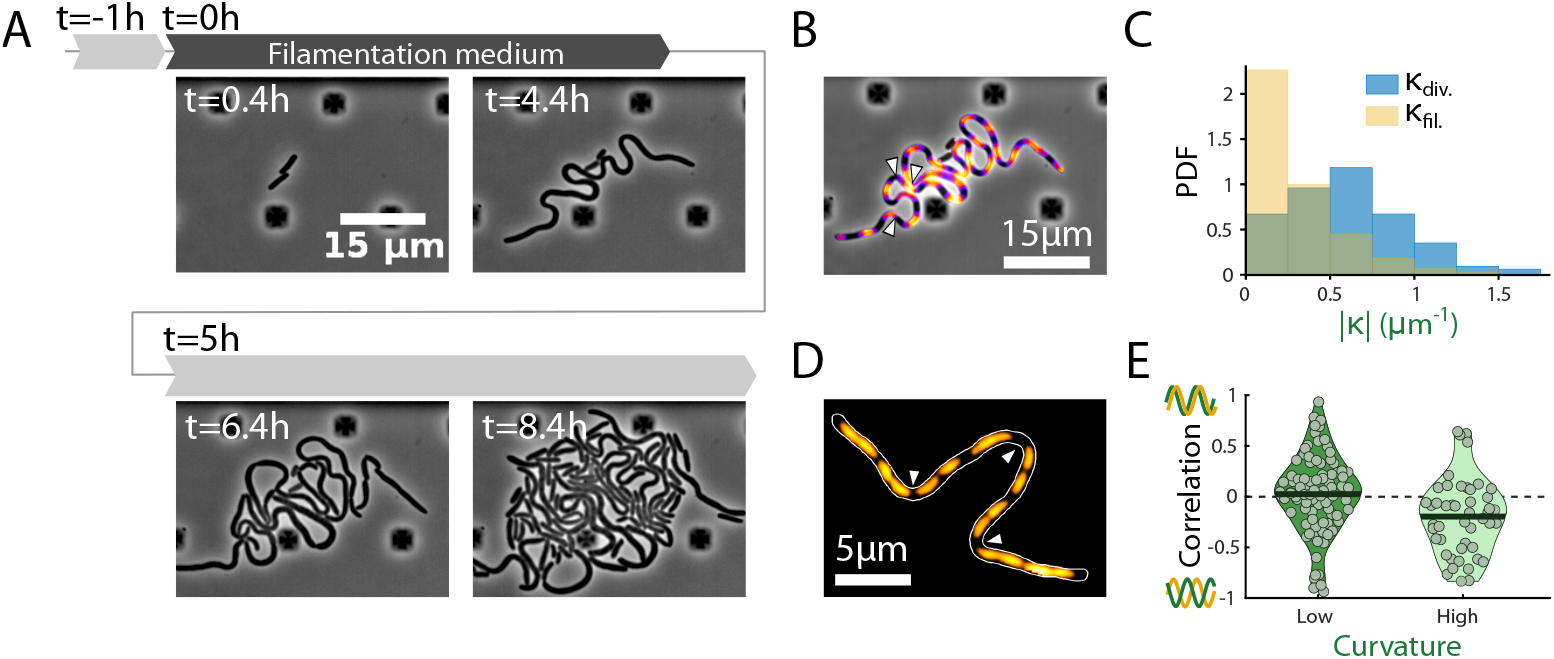
Min patterning and mechanical effects synergistically determine preferred division sites in post-stress conditions. **A**. Time-lapse of bacteria growing in a microfluidic chamber. After 1 h, aztreonam was added to the culture medium to induce filamentation. 5 h later, aztreonam was removed such that filaments resumed cytokinesis and the curvature at the division locations was calculated. **B**. 5 min average of phase contrast and fluorescent channels showing the location of MinD pulses just before the first division. The white arrows point at the 3 first divisions of the filament. **C**. Probability density functions of the curvatures for filaments at the division points (in blue) and for the whole filaments just before the corresponding divisions (in yellow). The means of the distributions are 0.6 ± 0.4 and 0.3 ± 0.3 *µ*m^−1^, respectively (sample size n=125). **D**. 5 min average of fluorescent channel showing DNA distribution avoiding curvature peaks (highlighted with white arrows) in a filament in the microfluidic device. **E** Violin plots showing the Pearson’s correlation coefficients between DNA signal and curvature. The correlation was calculated separating the buckled region (“high”) and the non buckled regions (“low”) as done with MinD in Fig. 2C (sample size n=47). Solid lines represent the means of the distributions (Low: 0.02; High: -0.20).

Under these conditions, we selected buckled filaments with a maximum curvature exceeding 0.5 *µ*m^−1^ and characterized the curvature at the division sites, |*κ*|_*div*._ (Fig. 5B). For the analysis, we included divisions in filaments longer than 8 *µ*m (either the originally selected ones or those resulting from divisions of longer filaments), ensuring that at least two division rings (or 3 sustained MinD pulses) were assembled in the cell (Fig. 5B). Fig. 5C shows the probability density function (PDF) of the curvature at the division points (blue), revealing that filament division is more likely to occur at sites with |*κ*|_*div*._ ≥ 0.5 *µ*m^−1^ (mean division curvature: ⟨|*κ*|_*div*._⟩ = (0.6 *±* 0.4) *µ*m^−1^). In contrast, the PDF of the curvature of filaments, |*κ*|_*fil*._, just before cytokinesis (Fig. 5 C, orange) shows a distribution shifted towards lower values of curvatures (⟨|*κ*|_*fil*._⟩= (0.3 ± 0.3) *µ*m^−1^). Further, given that the threshold curvature value for observing a negative correlation between curvature maxima and MinD sustained minima is 0.5 *µ*m^−1^, we calculated the fraction of divisions occurring above this threshold. Our results, revealed that 60% of divisions were above the threshold. If we normalize the curvature at the division by the mean curvature of the filament, the divisions above the mean curvature would account for 82% of the total.

The role of the Min oscillatory system in influencing chromosome positioning and segregation has been a topic of debate [54, 55]. In that regard, we additionally checked whether DNA segregation within the filaments followed the pattern dictated by MinD pulses, i.e., whether the DNA avoided the curvature peaks in filaments. To that end, we labeled and tracked the DNA in living filaments using SYTOX Orange (Methods). We found that the DNA was distributed in a discrete manner along the filament avoiding curvature peaks, Fig. 5D. This avoidance became more pronounced as curvature increased (see Fig. A6). These locations putatively mark the positioning of division septa when cytokinesis resumes [32, 53].To quantitatively assess the relationship between DNA distribution and the curvature of filaments we computed the correlation of their profiles (Fig. 5E). Our results revealed a significant anti-correlation (Mann-Whitney *U* test: *α* = 5·10^−2^ significance level). This demonstrates that curvature can influence chromosomal positioning, in line with previous studies showing that DNA can “sense” the curvature of the cell walls and adjust accordingly [54]. Consequently, these findings endorse the view that regions of high curvature, and thus mechanical strain, play a crucial, functional, role in determining preferential division sites in post-stress filaments.

## 3 Discussion

Previous investigations have studied *E. coli* filamentation and have established that Z-ring positioning is reallocated in response to the filament size [32]. Our study builds on this understanding by revealing how mechanical forces develop during *E. coli* filamentation and affect the Min system —the protein complex responsible for positioning the division ring. In particular, our findings offer a plausible explanation for the origin of the buckling phenomenon observed in filaments based on energy minimization principles: as filaments grow, elastic energy accumulates and is subsequently reduced through bending events. In this context, previous studies have shown that *E. coli* filaments exhibit elastic behavior when subjected to transient external forces. However, sustained external forces lead to significant cell wall synthesis, resulting in plastic deformations in which the localized activity of MreB plays a key role [19, 20, 56]. Additionally, hydrodynamic and adhesion forces have also been shown to influence filament shape [39]. This evidence suggests that a comprehensive quantitative description of the mechanical properties of filaments would require incorporating mechanisms and processes that were excluded from our model, as we deemed them unnecessary given the convincing results achieved using only a 1D linear elastic theory.

Regarding the impact of the buckling instability and curvature on Min signaling, we demonstrated that MinD can adjust in response to both naturally occurring curvatures and controlled curvatures induced by patterned growth substrates. Specifically, MinD accumulates in regions of increasing curvature, and for curvatures exceeding 0.5 *µ*m^−1^, our data indicated that MinD maxima avoid curvature peaks. Additionally, our signaling model showed that altering diffusion solely in the cytosol for the same diffusion drop did not affect the MinD pattern, while modifying diffusion exclusively in the membrane resulted in MinD accumulation, consistent with experimental observations (cf. Fig. 2B and Fig. 4B). In fact, previous studies suggested that increased protein binding leads to increased membrane occupancy, thereby reducing the diffusion of proteins across the membrane [57] and that curvature changes the binding affinity of membrane proteins affecting their diffusion [58]. This, together with our simulations, points to a decrease in membrane diffusion as the primarily responsible of the accumulation of MinD in regions of high curvature. Our FRAP experiments are consistent with this hypothesis. However, the observed diffusivity drop is not statistically significant, probably because in our experiments we cannot distinguish the fluorescence between membrane and cytosolic proteins. In that regard, it is worth mentioning that some studies have reported that cumulative light excitation slowed down GFP::MinD oscillations by about 10 s, potentially influencing FRAP results as well [59]. We also note that Min patterning modulation may be influenced by mechanisms beyond slower diffusion in buckled regions. For instance, mechanical stress has been shown to alter other biophysical properties, such as membrane potential, which in turn can impact divisome dynamics [12, 60–63].

Our study also explored the implications of our findings in post-stress conditions that lead to the resumption of cytokinesis in filaments. Given MinD’s role in inhibiting the divisome assembly in *E. coli*, and the observed tendency of Min to avoid curvature peaks, we hypothesized that Z-rings that assemble at these peaks would operate preferentially. Indeed, in our microfluidic experiments, where the stressor medium was removed —once the filaments reached high curvature values— the majority of division events occurred at curvatures exceeding the mean filament curvature. Furthermore, this observation aligned with our characterization of DNA distribution within the filaments, which also avoided curvature peaks. Ultimately, these results suggest the existence of a mechanical “memory” for cytokinesis, where the interplay between strain and signaling mechanisms defines the preferential division sites.

In a broader context, Min’s sensitivity to the shape of its “container” is not unique to *E. coli* [48, 58]. For example, in *B. subtilis*, an analogous complex to Fts, known as DivIVA, localizes to regions of highest curvature due to MinD’s preference for flatter membrane regions [64]. Mechanical deformations in minimal systems, such as liposomes containing Min proteins, have also been shown to alter Min patterning [65]. Specifically, MinE has been observed to distribute asymmetrically in response to liposome deformation. Interestingly, MinE accumulation and liposome deformation appear to mutually promote one another, suggesting a positive mechanochemical feedback loop that may play a role in our system.

Finally, while our study focuses on Min, other components of the divisome, such as FtsZ or membrane proteins such as MreB, could be affected by the mechanical stress in the membrane. In fact, a previous study has reported that MreB is correlated with the nucleoid position in *E. coli* unperturbed cells, but this correlation decreases when the cell geometry changes [66]. Thus, studying other components involved in the elongation, division, and the regulation of the mechanical properties of cells is crucial for the complete understanding of the relationship between signaling, filamentation, and cytokinesis and its implications for bacterial adaptation and antibiotic resistance [67].

## 4 Methods

### 4.1 Strain and growth conditions

Experiments were conducted using the *Escherichia coli* strain DH5*α* (NZYTech), transformed with the plasmid pYLS68 (P_lac_-YFP::minD minE::CFP), generously provided by Dr. Shih [68]. Overnight (O/N) cultures were initiated from a single colony and grown in LB medium supplemented with 100 *µ*g/ml ampicillin at 37°C with shaking at 220 rpm.

After overnight growth, cultures were diluted to a final OD_600_ of ∼0.01 in minimal M9 medium containing 0.4% glucose, 1 mM thiamine hydrochloride, 0.2% casamino acids, 100 *µ*g/ml ampicillin, and 100 *µ*M IPTG for plasmid induction. These morning cultures were incubated under the same conditions as the overnight cultures until they reached an OD_600_ of 0.2–0.3. Subsequently, they were further diluted to ∼0.005 to prepare the working cell culture.

### 4.2 Agarose pads and patterned growth substrates

Low melting point agarose pads were prepared following the established protocol [69] using M9-supplemented medium and 10 *µ*g/ml aztreonam to induce bacterial filamentation as previously described [39]. Subsequently, 2 *µ*l of the working culture were inoculated onto the agarose pads. The pads were incubated at 37°C for 15 min and then dried in a desiccator for 20 min. Once placed in the microscopy dish (IBIDI *µ*-Dish 35 mm, low), they were incubated at 37°C for 2 h to allow the cells to develop the filamentation phenotype.

To pattern the agarose growth substrates, master mold structures were fabricated on standard silicon samples. The fabrication process employed electron-beam direct writing on a Poly(methyl methacrylate) (PMMA) resist film. The electron-beam exposure, performed with a Jeol JBX-8100FS tool, was optimized to achieve the desired dimensions using an acceleration voltage of 100 keV and a beam current of 20 nA. After developing the PMMA resist using a conventional 1 : 3 MIBK/IPA developer, the resist patterns were transferred onto the silicon samples using an optimized Inductively Coupled Plasma-Reactive Ion Etching process with fluoride gases (SF_6_, C_4_F_8_). The remaining resist was removed with oxygen plasma.

The etched silicon patterns were used as molds for fabricating complementary patterns on PDMS. The silicon structures were silanized in gas with 1H, 1H, 2H, 2H-perfluorodecyltrichlorosilane (97%, stabilized with copper) for approximately 27 h by placing a droplet of the silanizer near the sample in a vacuum desiccator. A mixture of silicone elastomer base and curing agent (in a mass ratio of 1 : 10) was poured onto the silanized silicon pattern and cured in an oven at 90°C for 1 h. The cured PDMS stamp was cooled for 20 min at room temperature before being carefully peeled off from the silicon mold. This PDMS stamp contained a pattern complementary to that of the silicon mold.

To prepare the patterned agarose substrates, 2.5% low melting point agarose in culture medium with 1 *µ*g/ml aztreonam was poured onto the PDMS stamp and covered with a 2 × 2 cm coverglass. After refrigeration for 15 min, the coverglass was removed, and the PDMS was separated from the agarose. Finally, 2 *µ*l of cell culture with OD_600_ ≃ 0.3 was inoculated on the agar pad. The agar pads were dried in a laminar flow hood for 15 min before being transferred to a glass-bottom dish (IBIDI *µ*-Dish 35 mm, low). The same PDMS mold could be reused after proper cleaning: it was immersed in 100% ethanol, sonicated for 15 min, rinsed thoroughly with DI water, and dried in an oven at 60°C for 1 h.

The patterns were designed using the open-source mask design tool Nazca Design [70]. Four designs were created to control the curvature of the filaments (Fig. A3): three based on curved channels and one based on a lattice of pillars. The “wave” pattern featured a sequence of “s” curves formed by connecting three half-circular bends, each with a different radius of curvature (Fig. A3A). The “snake” pattern consisted of a sequence of “s” curves, where each “s” was formed by two half-circular bends, with the radius of curvature alternating every half rotation (Fig. A3B). For both patterns, the radius of curvature ranged from 0.35 *µ*m to 12 *µ*m. The “spiral” pattern was composed of half-circular bends with a radius of curvature increasing from 0.8 *µ*m to 10.4 *µ*m in increments of 0.8 *µ*m per half rotation (Fig. A3C). The width of the channels for these three patterns ranged from 0.7 *µ*m to 1.1 *µ*m. The fourth design was a triangular lattice of circles, each equidistant from its neighbors, with connected spaces in between. The diameters of the circles ranged from 2 *µ*m to 10 *µ*m, in 1 *µ*m increments, and the lattice constant ranged from 2.7 *µ*m to 10.7 *µ*m (Fig. A3D).

### 4.3 Microscope settings

Images were acquired using a Leica Thunder inverted microscope, equipped with a PL-APO 63 ×*/*1.40 oil objective and a CMOS camera (Leica DFC9000 GTC). Köhler illumination was applied to optimize image contrast. Excitation was performed using the Spectra-X Light Engine LED lamp, which features 8 solid-state LEDs, with the YFP filter settings (Ex: 495*/*517 nm, Em: 527*/*51 nm). The Leica Application Suite X (LAS X) software was used for microscope control and image acquisition. All experimental procedures were conducted at 37°C with temperature regulation provided by the IBIDI heating system.

Unless otherwise specified, time-lapses were acquired with an interframe time of 10 s over a 30-min duration, using 20% light source power, 30 ms exposure time, and 2 *×* 2 binning in the fluorescence channel.

### 4.4 Image and data analysis

Using Fiji [71], the background was removed from all images with a rolling ball radius of 50 pixels in the fluorescent channel. The brightness and contrast of the phase-contrast images were adjusted to maximize contrast based on the characteristics of each image. However, the levels of the fluorescence images were consistently adjusted between 200 and 5000 to ensure comparability. Segmentation was performed on the phase-contrast images using the WeKa plugin [72]. The resulting segmentation masks were then used to analyze both phase-contrast and fluorescence images with the MicrobeJ plugin [73], allowing us to extract quantitative data.

The coordinates of the medial axis obtained from MicrobeJ were analyzed to calculate the local curvature of the filaments using Mathematica [74] (Fig. 1A). The Mathematica code employs finite differences to estimate the first and second derivatives at each point of the medial axis of the filament. The output is a list of point pairs representing curvature and normalized length. To verify the curvature results, the known curvature of the agarose-patterned substrates was used as a control. To reduce noise, curvature data was smoothed using 5-min time-averages. Additionally, the curvature profile was further smoothed when the maximum filament curvature was *<* 0.3 *µ*m^−1^ to decrease noise. For filaments displaying buckling, this step was unnecessary as their curvatures typically reached ∼1 *µ*m^−1^, with a noise level of ∼0.1 *µ*m^−1^.

MinD intensity profiles obtained from MicrobeJ were analyzed to perform Fourier analysis of MinD oscillations in both space and time. These profiles were also used to compute the Pearson correlation between MinD intensity and curvature profiles using Mathematica. High- and low-curvature regions were defined using a curvature threshold of 0.5 *µ*m^−1^. High-curvature regions were defined from the left valley of the first peak where curvature exceeded 0.5 *µ*m^−1^ to the right valley of the last peak exceeding this threshold (Fig. A2C). Low-curvature regions were defined from the filament poles to these valleys. The correlation function was applied separately to each of these three regions (i.e., low curvature–high curvature–low curvature) for every filament.

To analyze the relationship between MinD intensity and filament curvature, MinD intensity profiles were processed to determine the MinD envelope (Fig. A2D). This approach avoided the influence of MinD pulse valleys. The maxima of MinD pulses were identified using the ‘FindPeaks’ function in Mathematica, followed by interpolation of these points and discretization to recover the original number of points. This process yielded a new MinD profile, representing the envelope of the original MinD signal. At each point along the filament’s arc length, *s*, the MinD intensity was normalized as:

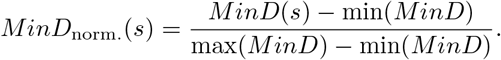

Curvature profiles were processed similarly, but without normalization. Curvature values were then grouped into the following bins: [0, 0.15), [0.15, 0.25), [0.25, 0.35), [0.35, 0.45), [0.45, 1), and [1, ∞). These bins, along with their corresponding *MinD*_norm._ values, were fitted to the function:

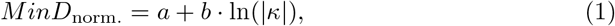

where *κ* is the curvature. The last bin was excluded from the fit as *MinD*_norm._ saturates and does not follow Eqn (1). From the fit, we obtained *a* = 0.69 ± 0.07 and *b* = 0.08 ± 0.04 with *R*^2^ = 0.9.

For agarose patterns, the last two bins were absent, as filaments did not reach those curvature values. Fitting the data to the same equation yielded *a*_pat._ = 1.00 *±* 0.98 and *b*_pat._ = 0.15 *±* 0.13 with *R*^2^ = 0.99.

### 4.5 FRAP experiments

The bacterial strain was grown and agarose pads were prepared as described in §4.1 and §4.2 respectively. Images were acquired using a ZEISS LSM 980 super-resolution confocal microscope equipped with a PLAN-Apochromat 63x/1.4 DIC M27-Oil lens, an Airyscan detector and ZEN software. All experiments were performed at 37 °C using the built-in incubation system.

Images were captured every 3 s for a total duration of 2.5 min. A 488 nm laser was used at 0.5% power with a 57 *µ*m pinhole. A master gain of 738 V and 304 V and digital offsets of −24116 and 0 were applied for the fluorescent and bright-field channels, respectively. Photobleaching was initiated after the fifth frame (15 s) by increasing the 488 nm laser power to 50% for 10 ms. The photobleached area was manually drawn using the polygonal selection tool to ensure it encompassed an entire MinD pulse. Recovery of fluorescence was monitored for a duration of 2.5 min, as photobleaching from repeated imaging precluded longer observations.

Photobleaching was performed on regular-length cells and filaments under various conditions. For regular-length cells, two scenarios were considered (Fig. A4): when MinD was fully concentrated at one bacterial pole, with no fluorescence at the opposite pole (Fig. A4 A), and when fluorescence was transitioning between the poles, with fluorescence visible at both poles (Fig. A4 B). For filaments showing three MinD pulses, photobleaching targeted the central pulse when MinD fluorescence was fully concentrated there, with no fluorescence at the poles (Fig. A4 C). For filaments showing more than three pulses, photobleaching was performed on a pulse located away from the poles (Fig. A4 D).

To assess MinD diffusion in filaments, photobleaching areas were drawn to cover approximately half of the filament. Filaments were categorized as either straight (maximum curvature *<* 0.5 *µ*m^−1^) or buckled (maximum curvature *>* 0.5 *µ*m^−1^). The coordinate origin was set at the edge of the photobleached area closest to the filament’s midpoint. In straight filaments, the analyzed pulse was the second pulse from this origin, whereas in buckled filaments, it was the first pulse located after the buckling region. Typically, this pulse was the second photobleached pulse, but depending on the filament’s geometry, it could also be the first or third pulse (Fig. 3 A & B, orange and blue dots).

The analysis of the selected pulse was conducted in Fiji by defining a circular region of interest (ROI) and plotting the Z-axis profile. Recovery time was defined as the first frame in which the fluorescence intensity of the photobleached pulse recovered to at least 30% of its pre-photobleaching intensity. The MinD oscillation period was measured by analyzing pulses in the non-photobleached regions of the filament and performing a Fourier transform to determine the oscillation period.

Recovery time (*t*_*r*_) was normalized by dividing it by the characteristic MinD oscillation period (*T*) for each filament, yielding a dimensionless recovery time. The distance (*d*) between the analyzed pulse and the first non-photobleached pulse (Fig. 3A, blue and orange dots) was measured using the segmented line tool in Fiji and normalized by the characteristic wavelength of each filament (*λ*). Recovery velocity was computed as the ratio of the dimensionless distance to the dimensionless recovery time, yielding a dimensionless velocity (*v*_*r*_*/c*). This methodology allowed for the comparison of MinD dynamics between straight and buckled filaments, highlighting how mechanical deformation influences protein diffusion and recovery behavior.

### 4.6 Microfluidic experiments

A microfluidic CellASIC ONIX2 BA04A03 system was used to study filament division after removal of the filamentation stressor. For each experiment, a single row of the device was utilized, pre-emptying wells 1, 2, 6, 7, and 8. *E. coli* cells were grown as described in §4.1, and 50 *µ*l of culture were transferred to well 8 of the microfluidic device. Well 2 was filled with 200 *µ*l of M9-supplemented media, while well 1 contained the same media supplemented with aztreonam at concentrations ranging from 1 to 10 *µ*g/ml.

To operate the device, a custom sequence was used: 1. Media from well 2 was flowed at 5 psi for 5 min to pre-clean the device. 2. Cells were loaded by flowing *E. coli* (well 8) at 2 psi. 3. After loading, media from well 2 was flowed at 1 psi for 30 min to 1 h to stabilize the cells. 4. Media containing aztreonam (well 1) was then flowed at 1 psi for 3 to 5 h. 5. Finally, the flow was switched back to aztreonam-free media from well 2 for 3 h to observe filament division points under the microscopy system described in §4.3. Division points were manually identified using the phase-contrast channel of microscopy time-lapses. Curvature at each division point was calculated for the frame immediately preceding division using the code described in §4.4. The curvature value was averaged over the three nearest points along the filament to the division point. The probability density function (PDF) of curvatures at division points was obtained using MATLAB [75] (‘histogram’ function using the ‘pdf’ normalization option). To compare this with the overall filament curvature, a second PDF was calculated for all curvatures of a filament. In this case, curvatures were smoothed using the MATLAB ‘smooth’ function. For this analysis, only filaments with a maximum curvature *>*0.5 *µ*m^−1^ and a minimum length of 8 *µ*m were considered.

To characterize DNA distribution within the filaments, the same microfluidic setup and loading protocol were followed. After cell loading, just media was flowed at 1 psi for 30 min. Subsequently: 1. Media containing aztreonam was flowed at 1 psi for 3 h. 2. Aztreonam media supplemented with SYTOX Orange (Invitrogen) was flowed for 30 min to 1 h to stain DNA. 3. Same media without SYTOX was then flowed at 5 psi for 5 min to remove excess background fluorescence. 4. Time-lapses were acquired for 5 min (10 s interframe time) using phase-contrast and fluorescence channels with the microscopy set-up described in §4.3 (SYTOX filter: Ex: 542*/*566 nm, Em: 578*/*610 nm). Correlation between SYTOX intensity and filament curvature profiles was calculated using a customized MATLAB script, following the same protocol applied for MinD (see §4.3). The poles of filaments (1 *µ*m from each end) were excluded from the analysis due to chromosome-free regions at the cell poles, consistent with previous findings [53].

### 4.7 Critical loading force

The same filaments used to analyze the DNA distribution in the microfluidic device were also employed to determine the critical buckling length. Time-lapse movies were acquired at 5 min intervals with an interframe time of 10 s. These time-lapses were projected into a single image that represents the average intensity of the signal of all frames using the ‘z project function’ in Fiji, using the average intensity option. Then, this projected image was thresholded to isolate the filaments, resulting in a binary mask. The MicrobeJ plugin was used to process this mask and obtain the medial axis, as described in §4.4.

Using the medial axis data, custom-written Mathematica code was employed to calculate the maximum curvature of each filament profile. These curvature values were then compared to the corresponding filament lengths, which were also obtained using MicrobeJ. The critical buckling load was calculated using the Euler formula:

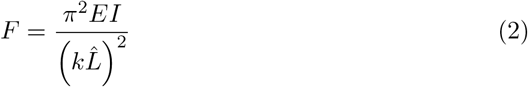

where *k* = 0.25 is the effective length factor for the filament [76]. The length of the filaments at which buckling started, 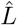, was chosen to be 10 *µ*m (see Fig. A1). *EI* represents the flexural rigidity of the filament, where *E* is the Young’s modulus and *I* is the second moment of inertia. The Young’s modulus, *E*, depends on the bacterial strain used. For the Dh5*α* strain used in our experiments, *E* is reported to range between 2 and 3 MPa [77]. For a thin hollow cylinder with radius *r* and wall thickness *b*, the second moment of inertia, *I*, can be expressed as *I* = *πr*^3^*b* [78]. Previous studies have reported a wall thickness of *b* = 4 nm for *E. coli* [19, 79]. Using a radius of *r* = 0.5 *µ*m [80], we calculate a flexural rigidity of *EI* = 4.7· 10^−21^ N*·* m^2^. Substituting these values into Eqn. (2), we obtain a critical loading force of *F* = 7.4 nN.

### 4.8 Mechanical model

*E. coli* filaments were modeled as a 2D chain of initial length *L*_0_ = 50 *µ*m, defined by *N* = 161 nodes. Each node, *i*, was represented by Cartesian coordinates 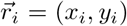 using a custom code written in *R* [81]. The characteristic distance between consecutive nodes was 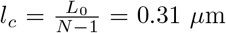. This resolution was chosen to ensure approximately six nodes per 0.3 *µ*m radius, corresponding to a maximum curvature of 3 *µ*m^−1^. The chain length *L* was selected to match the filament lengths observed in the experimental data. The initial coordinates for the chain were set as 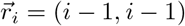, with *i* ranging from 0 to *N* − 1. A random perturbation was applied to the *y*_*i*_ coordinates, such that 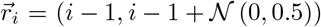, where 𝒩 (0, 0.5) denotes a normal distribution with 0 mean and 0.5 variance.

Two active forces were incorporated into the model: elastic and bending. For the elastic term, nodes were connected by linear springs with an elastic constant *K* and an equilibrium length *l*_0_ (*t*) that depends on time to simulate filament growth. The elastic energy for a node *i*, 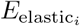, is given by:

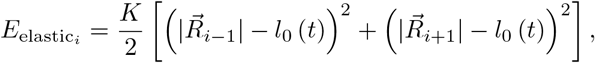

where 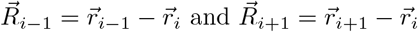.

The bending energy, which promotes filament flattening, depends on the angle formed by three consecutive nodes. Normalized vectors 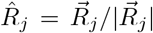 were used to ensure that the bending energy remains independent of the distances between nodes:

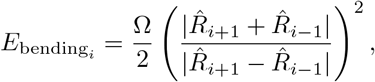

Thus, the bending energy reaches a minimum when the angle formed by three consecutive nodes is 180° and its maximum when the angle is 0°. To perform the simulations we implemented a dimensionless form of the energy, 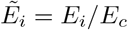, defined as a function of the characteristic energy 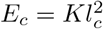:

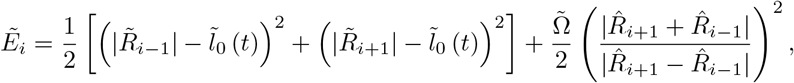

where 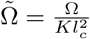

The equation for the dynamics of a node reads 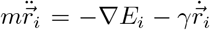, *m* being the “mass” of a node and *γ* a friction coefficient (energy dissipation). Disregarding the inertia term (low Reynolds number) we finally obtain that 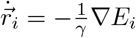 or, as a function of dimensionless units: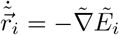 where a characteristic time *t*_*c*_ = *γ/K*, linked to the dissipation of the elastic energy, was defined. Filament growth was implemented by increasing the preferred length between nodes, *l*_0_ (*t*), as a function of time exponentially: *l*_0_ (*t*) = *l*_0_·2^*t/τ*^, *τ* being the doubling time. In our simulations the parameters shown in Table 1 were used.

**Table 1.**
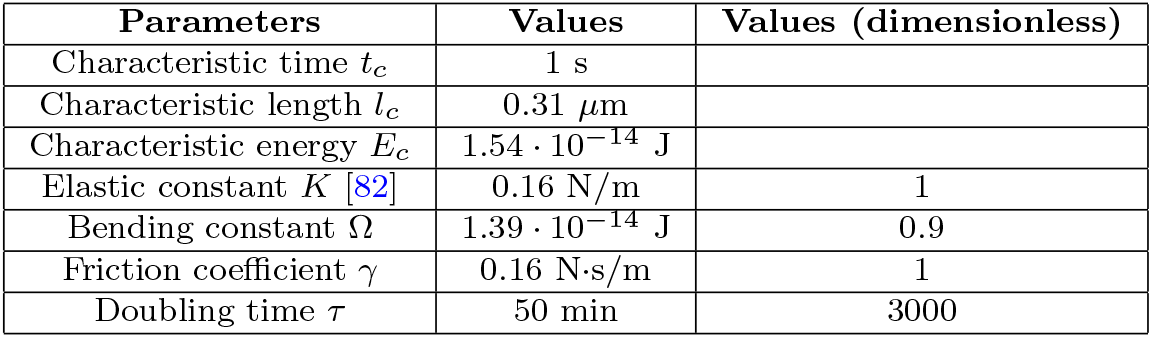
Table of parameters for the mechanical model.

| Parameters | Values | Values (dimensionless) |
| --- | --- | --- |
| Characteristic time $t_c$ | 1 s | |
| Characteristic length $l_c$ | 0.31 $\mu\text{m}$ | |
| Characteristic energy $E_c$ | $1.54 \cdot 10^{-14}$ J | |
| Elastic constant $K$ [82] | 0.16 N/m | 1 |
| Bending constant $\Omega$ | $1.39 \cdot 10^{-14}$ J | 0.9 |
| Friction coefficient $\gamma$ | 0.16 N·s/m | 1 |
| Doubling time $\tau$ | 50 min | 3000 |

The curvature at node *i* was defined as the inverse of the radius of the circle passing through nodes *i* − 1, *i*, and *i*+ 1, resulting in *N* − 2 curvature points. Statistics were generated by conducting 100 simulations with random seeds to initialize the perturbation of the *y*-coordinates. Simulation results were registered every 10 s to align with the experimental time-lapse interval. In simulations, to properly compared with the experiments, we implemented a pair-wise moving average that connects the middle points of adjacent spring connecting a node. That is, given a set of *M* coordinates of a filament, 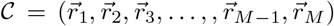 we generate the set of *M* coordinates 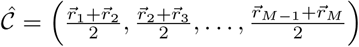.

### 4.9 Growth-induced elastic energy accumulation

Let us consider a chain of *N* (even) nodes as described in the previous section. For simplicity, the nodes are constrained to move along the *x* axis, and only elastic energy is considered. Initially, the nodes are symmetrically positioned with respect to *x* = 0, such that: 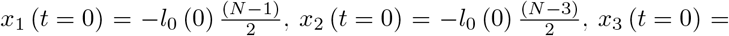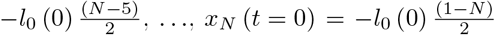. Consequently, 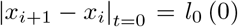, indicating that the system is initially in mechanical equilibrium. The dynamics of the nodes is then described by 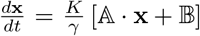 with **x** = (*x*_1_, *x*_2_, …, *x*_*N*−1_, *x*_*N*_)^*T*^ and 𝔸 and 𝔹 read,

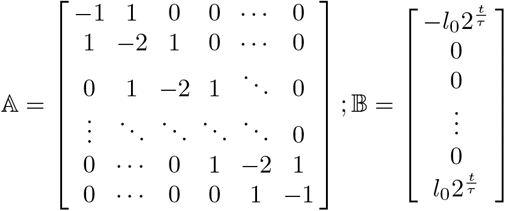

Due to the symmetry of the problem, the dimensionality of the system can be reduced to *N/*2, as *x*_1_ (*t*) = −*x*_*N*_ (*t*), *x*_2_ (*t*) = −*x*_*N*−1_ (*t*), and so on. This linear system can be solved, and it can be shown that |x_i+1_ − x_i_|_t_ > 0 ∀ *t*. As expected, the exponential growth of the equilibrium length, 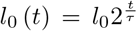, leads to increasing separation of the nodes over time.

Furthermore, elastic energy accumulates in the central springs compared to the springs at the ends of the chain. For instance, in the simplest case of *N* = 4 (one central spring surrounded by two springs at the ends), the ratio of the lengths between the end spring and the central spring over short times is given by:

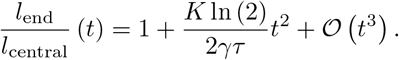

This implies that, regardless of the spring constant *K*, the dissipation coefficient *γ*, or the doubling time *τ*, elastic energy preferentially accumulates in the central spring due to growth.

### 4.10 Min signaling model

The signaling model developed by Meinhardt and J. de Boer [52] was used for the simulations presented in Fig. 4. The model considered two proteins that can be in two different states: membrane-bound *MinD*_*mem*._ and *MinE*_*mem*._, and cytoplasmic *MinD*_*cyto*._ and *MinE*_*cyto*._. Parameter values from the original study were used, with modifications to the characteristic time and length scales to match our experimental observations: approximately 20 MinD pulses over 30 min. Specifically, the characteristic time was set to *t*_*c*_ = 2.4·10^−3^ s, and the characteristic length to *l*_*c*_ = 0.29 *µ*m.

In the model, *E. coli* filaments were represented as a 2D chain of initial length *L*_0_ = 50 *µ*m with *N* = 172 nodes (*l*_*c*_ = 1 in dimensionless units). Protein concentrations were defined at the nodes. Filaments grew exponentially over 30 min, with a doubling time *τ* = 50 min: *L*(*t*) = *L*_0_2^*t/τ*^. In the simulation the length of the filaments was increased in discrete steps such that two new nodes were added at the ends of the filaments when the length increased by 2 × *l*_*c*_. The partial differential equations were numerically integrated using the FTCS algorithm (null flux boundary conditions) [83].

We also performed controls to evaluate the effect of the reduction of cytoplasmic diffusion (Fig. A5B). Flat filaments (|*κ*| *<* 0.1 *µ*m^−1^) retained 100% of the value of the diffusion coefficients. To calibrate the relationship between diffusion and curvature, the experimental dependence of MinD intensity as a function |*κ*|, Eqn. (1), was used (see Fig. 2D):

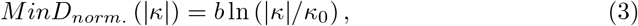

where *κ*_0_ = *e*^−*a/b*^ = 1.78*·*10^−4^ *µ*m^−1^. Using this relationship we mapped the simulated MinD intensity, *MinD*_*sim*._ = *MinD*_*mem*._ + *MinD*_*cyto*._, to values of the mimicked curvature |*κ*|_*sim*._:

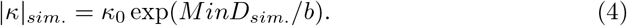

To calibrate the relationship between a drop in diffusion coefficient and the curvature, we ran simulations using membrane diffusion coefficients of 100%, 70%, 60%, 50%, and 40% in a region 1.5 *µ*m-wide, and obtain the corresponding *MinD*_*sim*._ values and |*κ*|_*sim*._. We subsequently mapped diffusion and curvature values using a logarithmic dependence [46]:

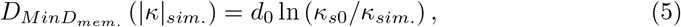

resulting in *d*_0_ = 0.17 *µ*m^2^/s and *κ*_*s*0_ = 7.73 *µ*m^−1^.

For the simulation shown in Fig. 4C, 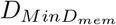 and 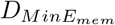 were reduced to 40% of their values (*s* = 1.5 *µ*m band). Thus, using Eqn. (5), |*κ*|_*sim*._ = 1.27 *µ*m^−1^, and the simulated angle of the bending was *θ* = *s* |*κ*|_*sim*._ ≃ 109°. The diffusion drop was applied after 10 min of cellular growth.

## Acknowledgements

We thank Dr. Shih for providing us with the pYLS68 plasmid and Prof. Ulrich Schwarz for his valuable comments during the preparation of this manuscript. We are also grateful to Enrique Navarro Raga from SCSIE at UV for his assistance with the FRAP assays and to Amadeu Griol from NTC for his support with the fabrication methodology.

I.G. acknowledges funding from the Margarita Salas Fellowship (Ref. MS21-128) and the HORIZON-MSCA-2022-PF-01-01 program (grant no. 101107228). I.D., J.H., and A.M. acknowledge support from the European Commission through the CHI-RALFORCE Pathfinder project (grant no. 101046961) and from the Conselleria de Educacin, Universidades y Empleo under the NIRVANA grant (PROMETEO Program, CIPROM/2022/14). J.B. acknowledges support from grant PID2022-137436NB-I00 and the research network RED2022-134573-T funded by ‘Ministerio de Ciencia e Innovación’ (MCIN/AEI/10.13039/501100011033) and by ERDF ‘A way of making Europe’ by the E.U. Additional support to J.B. was provided by the E.U. COST action CA22153 ‘European Curvature and Biology Network’ (EuroCurvoBioNet).

## Declarations

- Funding: See acknowledgments.
- Conflict of interest/Competing interests: The authors declare no competing interests.
- Ethics approval and consent to participate: ‘Not applicable’.
- Consent for publication: ‘Not applicable’.
- Data availability: Data for this study will be available upon request.
- Materials availability: ‘Not applicable’.
- Code availability: Codes for this study will be available upon request.
- Author contribution: M.N.: experiments, data analysis, writing, reviewing and editing. L.G.: numerical simulations (mechanical and signaling models). I.D.: micro-fabrication and writing (methods). J.H. and A.M.: micro-fabrication. I.G.: experiments, data analysis, writing, reviewing, editing, conceptualization and supervision. J.B.: writing, reviewing, editing, data analysis, conceptualization, funding acquisition, and project supervision.

## Appendix A Extended Data

**Fig. A1.**
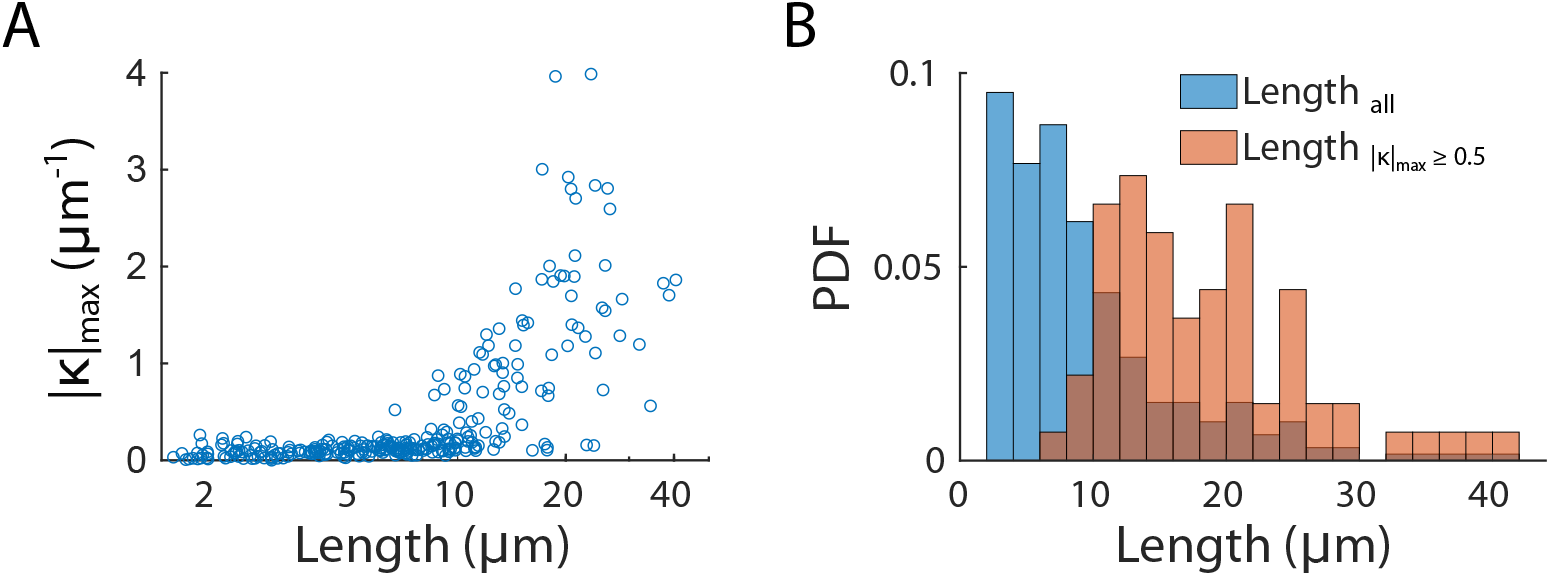
Critical length for buckling onset. **A**. Maximum curvature |*κ*|_*max*_ presented in 300 filaments of different lengths. |*κ*|_*max*_ stays below 0.5 *µ*m^−1^ up to a length of around 10 *µ*m. **B**. Probability density functions (PDF) for the lengths of 300 analyzed filaments (blue) and the length of the filaments whose |*κ*|_*max*_ ≥ 0.5 *µ*m^−1^.

**Fig. A2.**
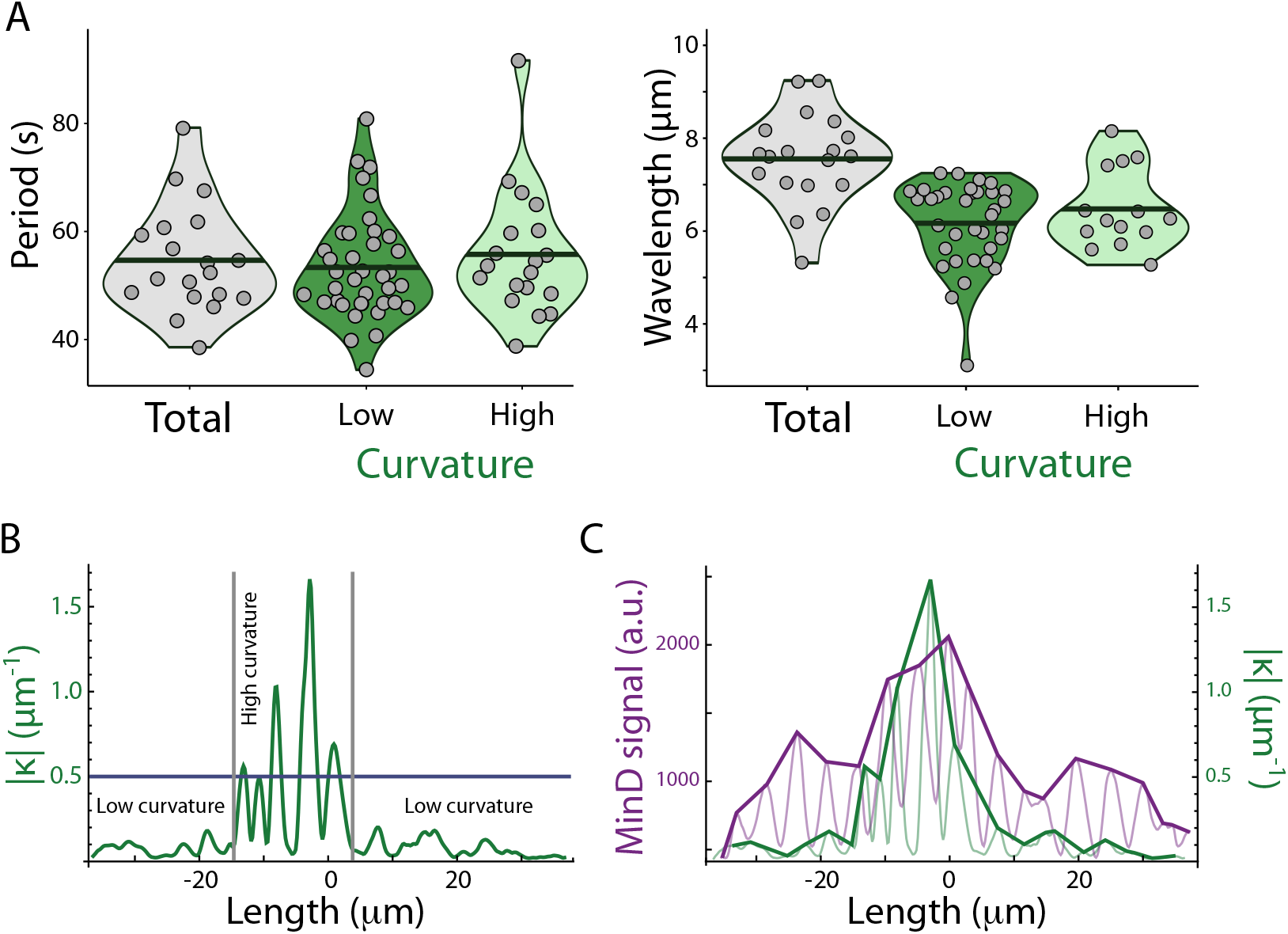
Experimental characterization of the wave properties of MinD patterns and curvature. **A**. Violin plots of Fourier analysis showing the distribution of values of period *T* (left), and the wavelength *λ* (right) (sample size: *n* = 19). In “Total”, these quantities were calculated for the whole filaments. In “Low” and “High”, *T* and *λ* were analyzed in the non-buckled and buckled regions separately, as indicated in **B**; for *λ* only the fragments whose length ≥ 8 *µ*m were considered. Black solid lines represent the mean (Period Total: 55 s, Low: 53 s, High: 56 s; Wavelength Total:7.6 *µ*m, Low: 6.3 *µ*m, High: 6.5 *µ*m). **B**. Curvature profile of an illustrative filament showing how low and high curvature regions are defined using a curvature threshold |*κ*| = 0.5 *µ*m^−1^. **C**. Example of envelope curves of MinD and curvature profiles of an illustrative filament used to analyze the effect of the MinD accumulation around peaks of high curvature.

**Fig. A3.**
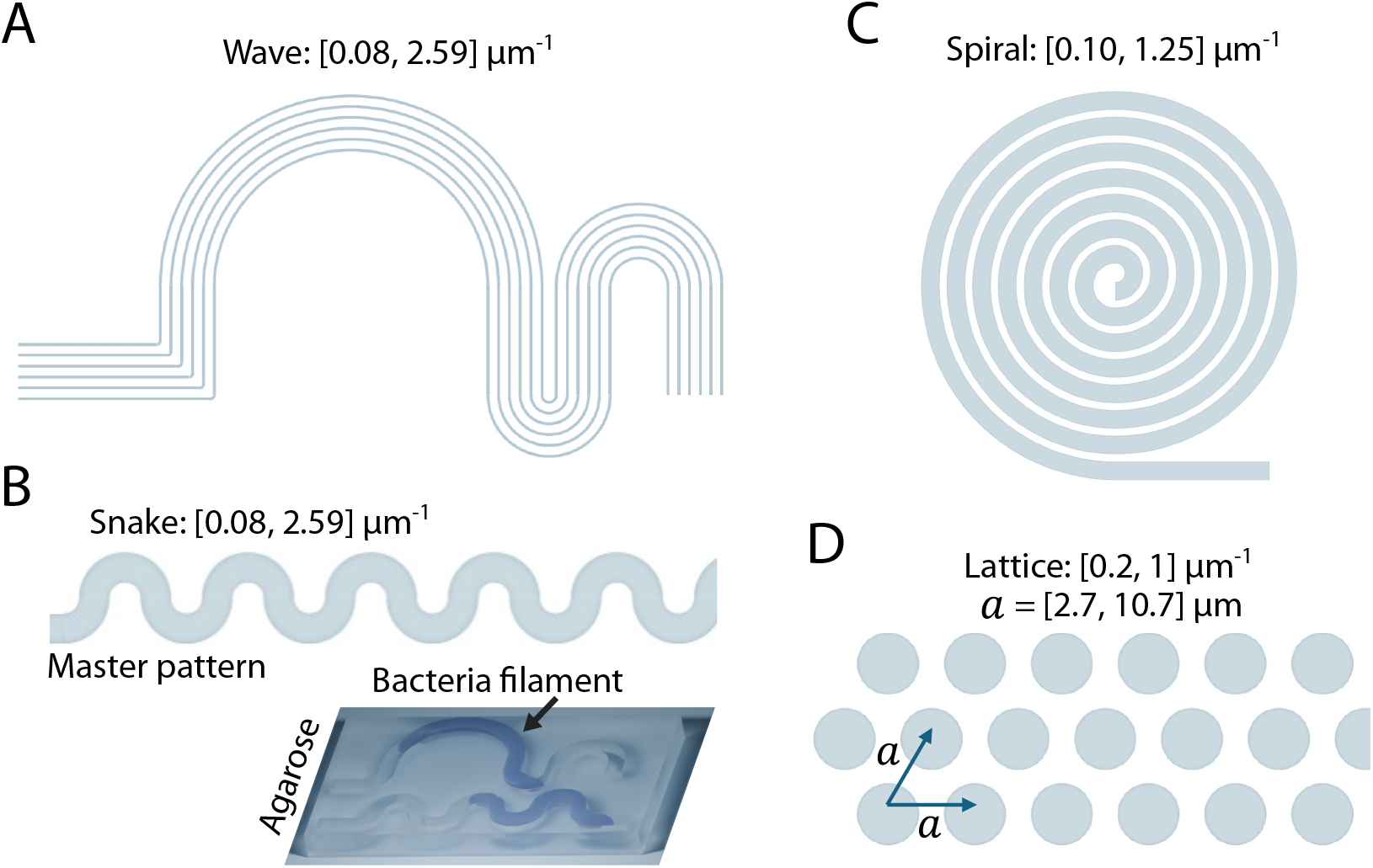
Micro-pattern designs for growing substrates. **A**. The “wave”: a sequence of “s” curves formed by connecting three different half circular bends (each with a different radius of curvature). **B**. The “snake”: a sequence of “s” curves, where each “s” shape is formed by two half circular bends, with its radius of curvature changing sign every half rotation. For these two first patterns the radius of curvature ranged from 0.35 *µ*m to 12 *µ*m. Cartoon of the resulting patterned agarose growth substrate. **C**. The “spiral”: a sequence of half-circular bends whose radius of curvature goes from 0.8 *µ*m to 10.4 *µ*m increasing 0.8 *µ*m every half rotation.**D**. The “lattice”: a triangular arrangement of circles where each circle is equidistant from its nearest neighbors, with connected space between them. The diameters of the circles ranged from 2 *µ*m to 10 *µ*m, and the lattice constant *a* ranged from 2.7 *µ*m to 10.7 *µ*m.

**Fig. A4.**
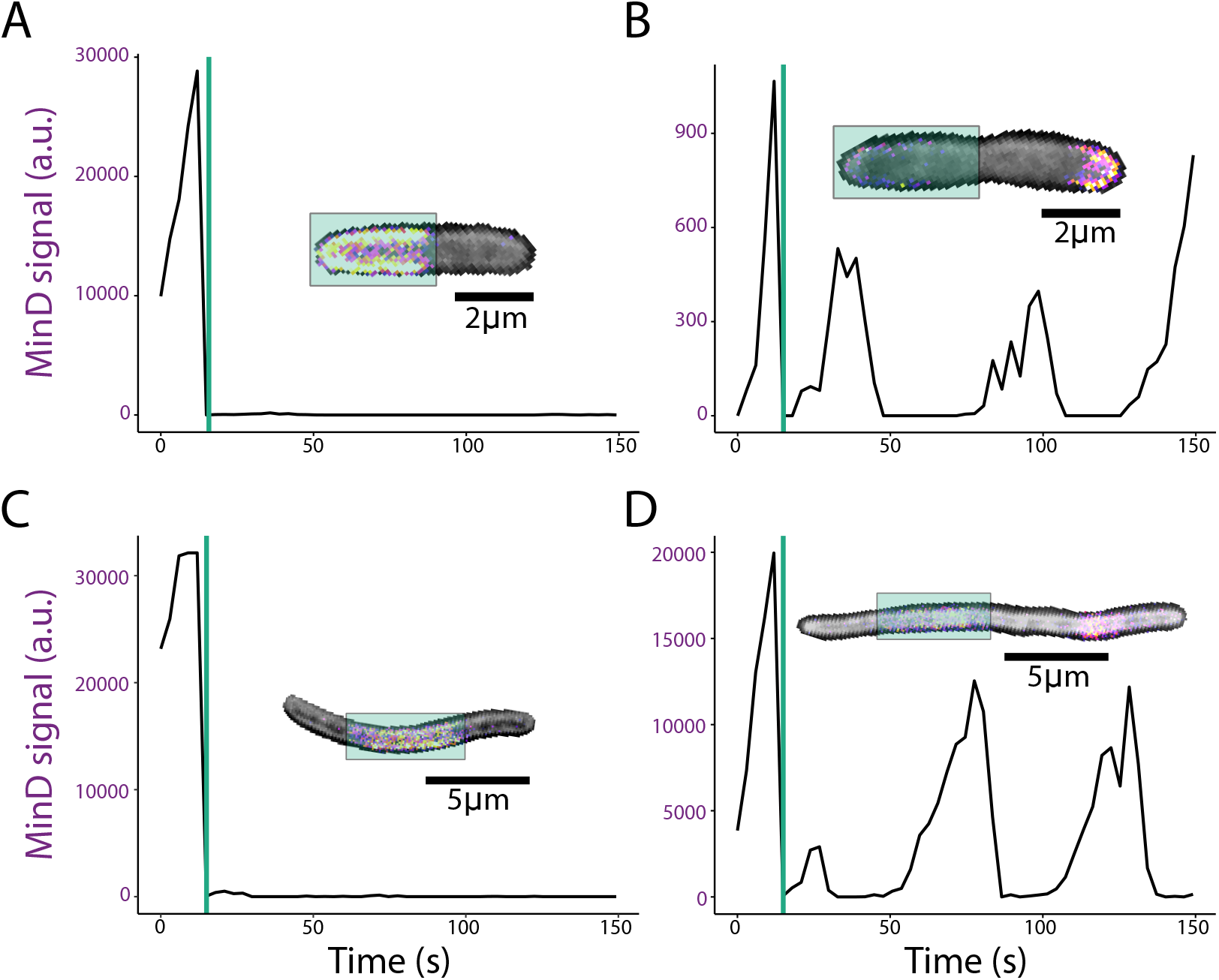
Control FRAP experiments. Time evolution of MinD intensity in the analyzed pulse (receiver). Vertical lines indicate photobleaching events. Insets show the composite of bright-field and MinD fluorescence channels of the analyzed bacterium. The colored area highlights the photobleached region. **A**. Photobleaching of a MinD pulse when all the MinD protein in the cell is concentrated in the photobleached area in a bacterium. This resulted in no MinD recovery for the time studied. **B**. Photobleaching of a MinD pulse when MinD is present in both the photobleached area and outside of it. This resulted in a recovery of MinD with its characteristic oscillation period. **C**. Filament with three MinD pulses, poles pulses in phase and middle pulse in counter-phase. The middle pulse is completely photobleached when all MinD is accumulated there. This resulted in no MinD recovery for the time studied. **D**. A filament with more than three pulses always has at least two pulses in phase. Photobleaching one pulse results in a fluorescence recovery.

**Fig. A5.**
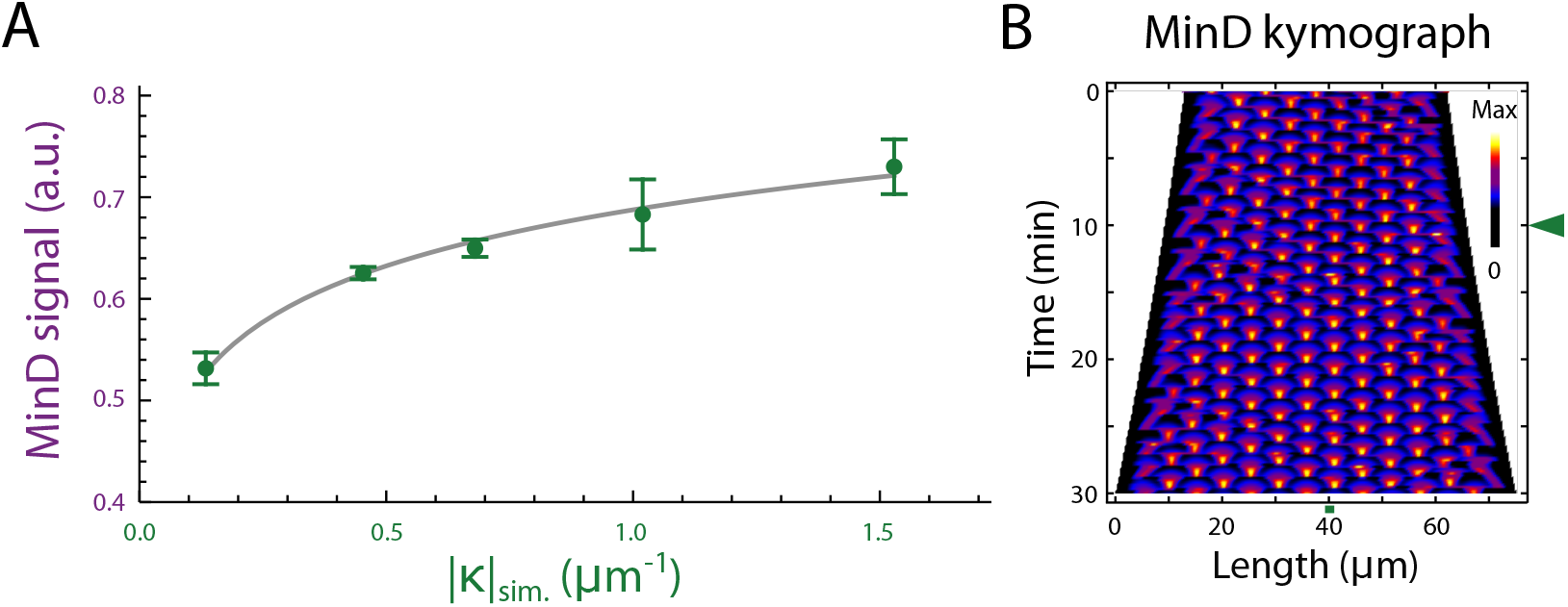
Signaling model: calibration and effect on MinD patterning when changing the cytoplasmic diffusion. **A**. Normalized simulated MinD intensity, *MinD*_*sim*._, calibrated as a function of the simulated membrane curvature (|*κ*|_*sim*._). Solid gray line: fitting of the experimental data to Eqn. (1). Green circles: simulation data (*n* = 5) for different 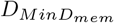 and 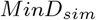. diffusion coefficients (100%, 70%, 60%, 50%, and 40% of their values). **B**. Kymograph of *MinD*_*sim*._ intensity when the diffusion coefficients of 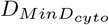 and 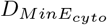 are 40% of their regular values. Changes on the coefficients are applied at *t* = 10 min (green arrow) onward in a 1.5 *µ*m band (horizontal green band).

**Fig. A6.**
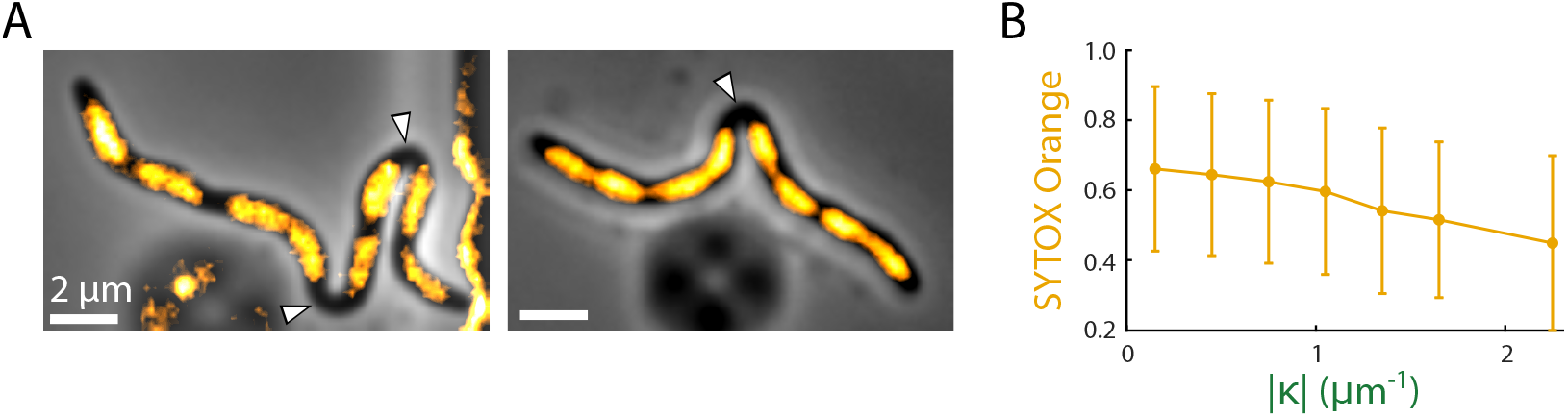
DNA distribution in buckled filaments. **A**. Merged images (phase contrast + DNA fluorescent signal) in illustrative filaments undergoing buckling. The DNA avoids the curvature peak (white arrows). **B**. Normalized SYTOX signal (DNA) as a function of the curvature. Circles indicate the mean intensity value (bars: standard deviation; *n* = 47 filaments). Data were binned in the following intervals of |*κ*| (*µ*m^−1^): [0, 0.3), [0.3, 0.6), [0.6, 0.9), [0.9, 1.2), [1.2, 1.5), [1.5, 1.8), and [1.8, ∞).

## Appendix B Movie Captions

- **Movie 1**. Phase contrast time-lapse of a filament growing on an agarose pad (10 *µ*g/ml aztreonam). Interframe time: 10 s.
- **Movie 2**. Simulation of a filament (mechanical model) undergoing growth-induced buckling. Bending and elastic energies are shown by means of color scales.
- **Movie 3**. Fluorescence channel (MinD::YFP, false color) of the filament shown in Movie 1.
- **Movie 4**. Filament growing in a ‘spiral’ micro-patterned agarose pad (1 *µ*g/ml aztreonam): phase contrast + fluorescent channel (MinD::YFP, false color). Interframe time: 1 min.
- **Movie 5**. Filaments in a microfluidic device resuming cytokinesis after aztreonam is removed from the medium. Interframe time: 1 min.

